# Antimicrobial Activity of Tetrabromobisphenol A (TBBPA) against *Staphylococcus aureus* Skin Infections

**DOI:** 10.1101/334193

**Authors:** Chang Wang, Fang Ji, Fengjie Chen, Bolei Chen, Zhen Zhou, Zhi Li, Yong Liang

**Affiliations:** Institute of Environment and Health, Jianghan University, Wuhan, P.R. China; School of Medicine, Jianghan University, Wuhan, P.R. China; Research Center for Eco-Environmental Sciences, Chinese Academy of Sciences, Beijin; Institute for Interdisciplinary Research, Jianghan University, Wuhan, P.R. China; Key Laboratory of Optoelectronic Chemical Materials and Devices, Ministry of Education, School of Chemical and Environmental Engineering, Jianghan University, Wuhan, P.R. China; Department of Orthopedis, Wuhan General Hospital of Guangzhou Command, 627 Wuluo Road, Wuhan, P. R. China

**Author notes:** Address correspondence to Yong Liang. Chang Wang and Fang Ji are co-first authors.

## Abstract

Tetrabromobisphenol A (TBBPA) is a brominated flame retardant with selective antimicrobial activity against Gram-positive bacteria. We show that TBBPA exerts bactericidal effects by damaging the cell wall and membrane of *Staphylococcus aureus* (SA) without inducing antimicrobial resistance. *In vivo* skin infection assays indicate that a low dose of TBBPA could contribute to wound closure and attenuate SA infection and inflammatory infiltration. TBBPA has potential for use as an antimicrobial agent against Gram-positive pathogens.

---

*Staphylococcus aureus* (SA) remains an important cause of cross infections, including uncomplicated skin infections and severe fatal infections such as pneumonia, osteomyelitis and endocarditis. Bacterial infections caused by resistant pathogenic microorganisms are commonly found in hospitals worldwide (1-3). Increasing development of antimicrobial resistance is a global concern; vancomycin-resistant SA raises the possibility of incurable infection (4, 5). Also, abuse of antibiotics has become a widespread health problem in recent years. There is an urgent need to develop antibacterial agents, based on antimicrobial peptides, plant products or synthetic drugs that are highly efficient and have a low incidence of resistance. Tetrabromobisphenol A (TBBPA) is the most widely used brominated flame retardant, with an estimated production of 200,000 tons per year. TBBPA is used in epoxy resins to retard fire (6). Previous findings from our laboratory showing Gram-positive specific antimicrobial activity of TBBPA (7) led us to further study this activity against common pathogens, focusing on *in vitro* and *in vivo* antibacterial activity against SA. The primary objective of this study was to evaluate the use of TBBPA as a treatment for bacterial infection.

In the susceptibility studies, the common pathogens *Enterococcus faecium, Staphylococcus aureus, Klebsiella pneumoniae, Acinetobacter baumanii, Pseudomonas aeruginosa* and *Escherichia coli* were tested. TBBPA selectively exerted antimicrobial effects against Gram-positive bacteria, consistent with our previous findings (7). TBBPA inhibited the *in vitro* growth of SA and MRSA at a concentration of 4 or 2 µg/mL, respectively (Table 1). These common strains are known as the “ESKAPE” pathogens, which cause the majority of hospital infections in the US, with MRSA infection leading to identifiable skin and soft-tissue infections in US hospitals (8, 9). These results indicate that TBBPA is a valuable antibacterial agent, with high specific activity against Gram-positive pathogens.

**Table 1:** Minimal inhibition concentrations (MICs) of TBBPA on the typical pathogens.

| Strain | MIC (µg/mL) |
| --- | --- |
| **Gram-positive bacteria** |  |
| *Staphylococcus aureus* 25923 | 2 |
| MR - *Staphylococcus aureus* | 4 |
| *Enterococcus faecium* | 4 |
| **Gram-negative bacteria** |  |
| *Klebsiella pneumoniae* | > 128 |
| *Acinetobacter baumannii* | > 128 |
| *Pseudomonas aeruginosa* | > 128 |
| *Escherichia coli* DH5α | > 128 |

We evaluated the development of antimicrobial resistance to TBBPA in SA. SA was cultured for 40 days with daily transfers in the presence of antibiotics (concentrations were constant or increased stepwise). The minimum inhibitory concentrations (MICs) of ampicillin (AMP) and chloramphenicol (CHL) against SA increased rapidly and AMP- or CHL-resistant SA was established with a MIC increase of 25- or 12-fold, respectively. In contrast, we were unable to obtain TBBPA-resistant SA in the presence of a low concentration (4×MIC) of TBBPA. Serial 60 passage of SA in the presence of sub-level MIC of TBBPA for 40 days failed to produce highly resistant strains (Fig. 1), indicating a lower development of resistance to TBBPA compared with AMP and CHL.

**Figure 1:**
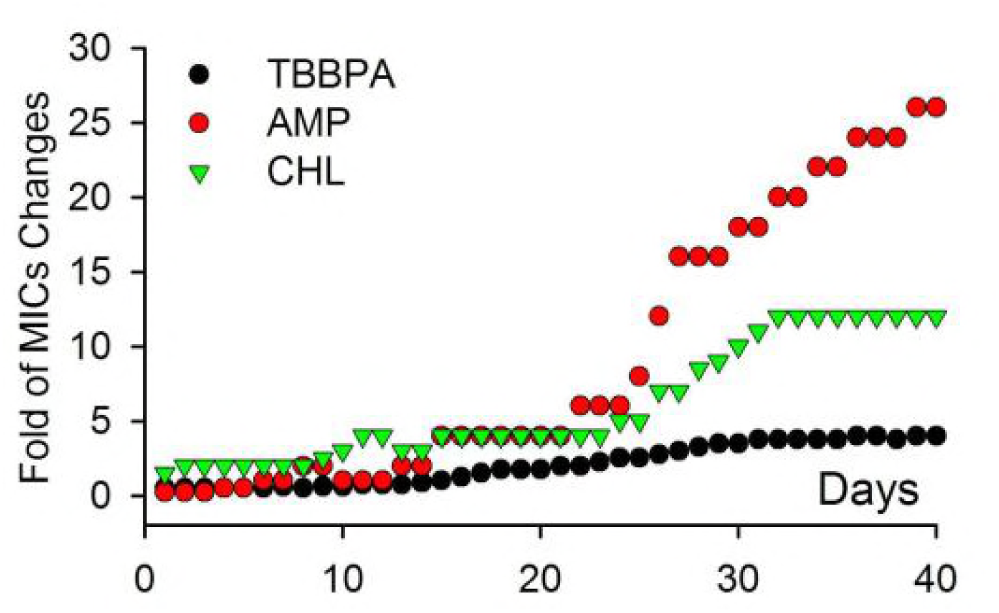
Antimicrobial resistant characterization of SA to TBBPA, AMP (ampicillin), and CHL (chloramphenicol) for 40 days. The results shown are representative of experiments performed at least three times.

Gram-positive and Gram-negative bacteria have distinct differences in cell wall structure, leading to the possibility that the specific antimicrobial activity of TBBPA may relate to the presence of teichoic acid in the peptidoglycan layer. This led us to try to understand the mechanism of antimicrobial activity by analyzing changes in cell wall morphology using scanning and transmission electron microscopy (SEM and TEM). TBBPA exposure resulted in loss of normal cell morphology with significant cracks in the cell envelope (Fig. 2.2). Control cells had an intact cell envelope with protoplasts inside (Fig. 2.1, 2.3). TBBPA-treated cells showed significant protoplast leakage through the cell envelope (Fig. 2.4), formation of a mesosome-like structure (Fig. 2.5) and cellular content leaked through cracks in the cell envelope (Fig. 2.6). Damage to the cell wall eventually led to the death of TBBPA-treated cells. The formation of a mesosome-like structure indicated alteration of the cell membrane. It has been reported that some antimicrobial peptides and synthetic antibiotics induced a mesosome-like structure during cell division (10). These results demonstrated that TBBPA may exhibit antimicrobial effects by inducing cell wall damage, affecting cell membrane stability.

**Figure 2:**
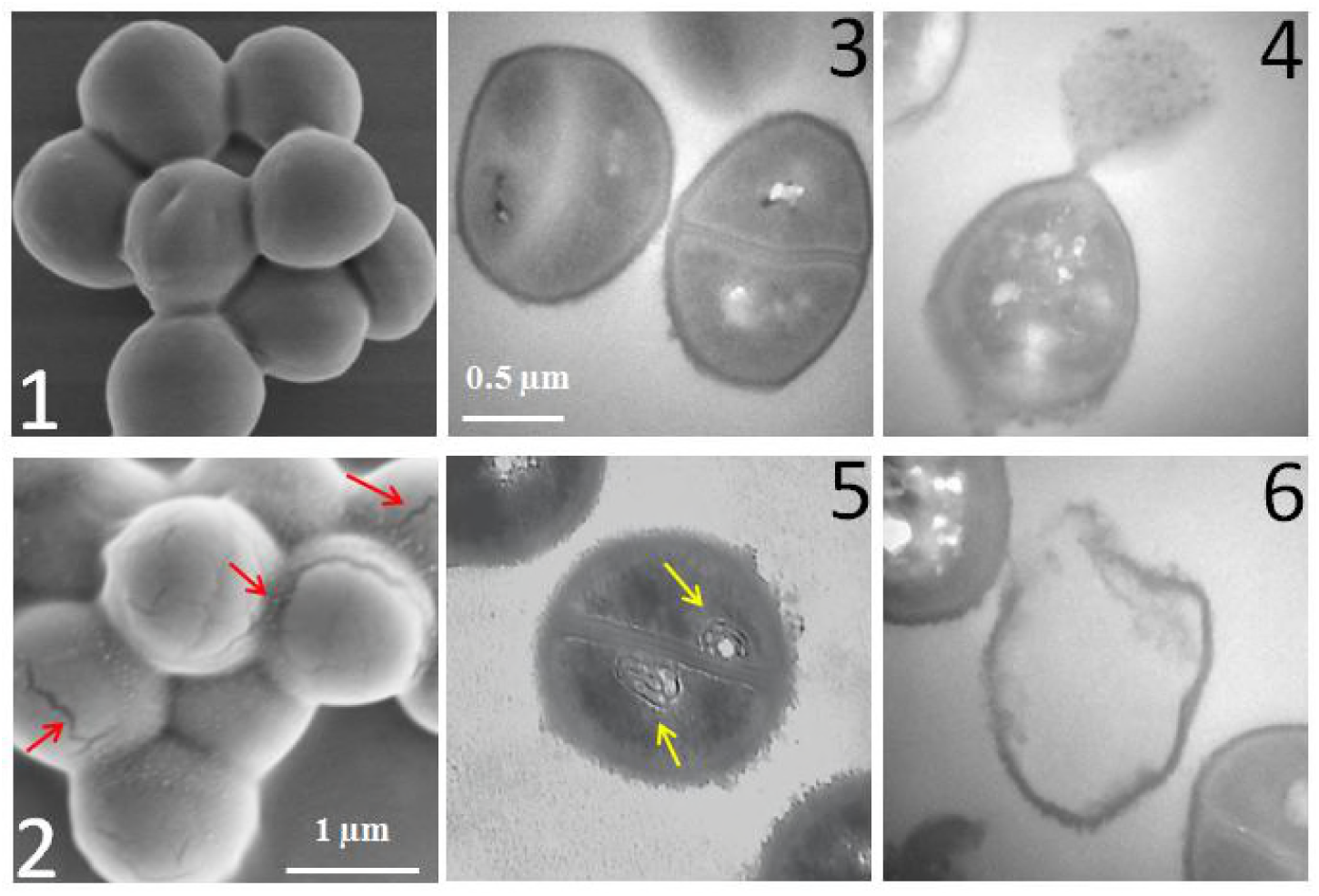
TEM and SEM images of TBBPA-exposed SA. The culture of SA with an OD600 of approximately 0.6-0.8 was treated with 20µg/ml of TBBPA (2, 4, 5 and 6) or an equivalent amount of DMSO as control (1, 3) for 1 h. The treated cells were studied using TEM (1, 2) and SEM (3-6). Red arrows indicate the small crevices on the cell wall (2) and yellow arrows indicate the mesosome-like structure (5). The severely damaged cell wall (2, 4, and 6), the protoplast leaking out (4), and the formation of mesosome-like structure (5) are observed. The scale bars (The same for 1 and 2; 3 and 4-6)

Therapeutics against bacterial infections are limited and development of novel therapeutics to treat drug-resistant infections has been a concern, as many pathogens are resistant to most currently available antibacterial agents. Therefore, the development of efficient antimicrobial agents with a low incidence of resistance is urgently needed. Animal models of skin infection are important tools to elucidate the potential of new antimicrobials to prevent or reduce the severity of infection (11). We investigated whether TBBPA was effective as an antimicrobial against SA using an *in vivo* skin infection model. We assessed the effect of TBBPA on SA-induced pathological changes in the skin and determined the number of viable SA cells in skin wounds. Wounds were excised on day 11 and the number of viable SA cells present in the tissue was determined (Fig. 3B). There were significant differences in viable SA cell numbers isolated from fester and exudates in the wound tissue. Viable SA cell numbers were significantly reduced when mice were treated with TBBPA or AMP, compared with untreated mice. AMP and TBBPA treatment significantly reduced bacterial load, which was measured using the plate count method (Fig. 3F). Wound closure of vehicle control did not change over the 7 day test period. Treatment with TBBPA resulted in obvious wound recovery (Fig. 3A and B) and significantly reduced the CFUs of SA, similar to the outcome of AMP treatment. Thus, TBBPA could be used as a treatment for SA-induced skin infections. Histological examination of the wound tissue revealed neutrophil infiltration and epithelialization in different treatment groups. Treatment with TBBPA resulted in a significant thinning of the epidermis layer and less dissociated epithelial cells in dermis layer (Fig. 4). The increase in epithelialization suggests that the recovery process may have been delayed. Untreated infections showed significant epithelialization on day 11, having a thicker epithelial layer compared with normal skin tissue. TBBPA demonstrated good *in vivo* activity in a mouse skin infection model which was comparable to antibiotics in some cases; the inflammatory response of the epidermis and muscle cells caused by SA was significantly reduced by TBBPA treatment. Overall, these results were consistent with the hypothesis that TBBPA exhibits *in vivo* antimicrobial activity and provides protection against SA skin infection.

**Figure 3:**
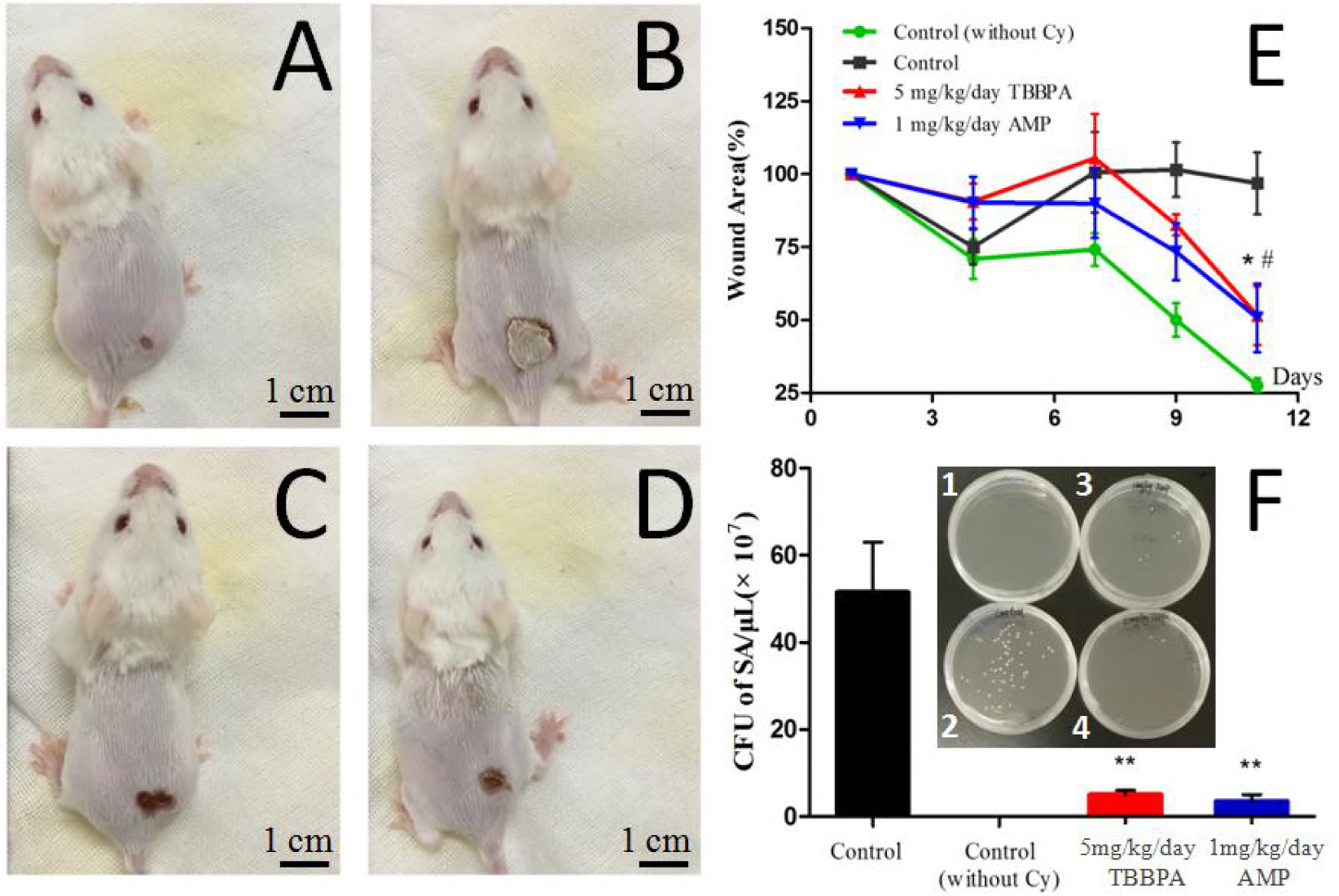
Skin SA-infection model to study the antimicrobial activity of TBBPA in BALC/c mice. Mouse was treated with 60 mg/kg cyclophosphamide (Cy) for 3 days and maintained for one more day following subcutaneous inoculation with SA on day 1. All the mice in group B-D were pretreated with Cy to normalize the individual variations and suppress the immune system. A, SA-infection without Cy treatment (Control without Cy); B, SA-infection (Control); C, SA-infection and 5 mg/kg/day TBBPA treatment (5 mg/kg/day TBBPA); D, SA-infection with 1 mg/kg/day Ampicillin treatment (1 mg/kg/day AMP); E, Relative wound area of necrotic ulcer shown against days after infection, # and * indicate the significant different compared to reference group (Control without Cy, n=6,*P<*0.01); F, SA bacteria cultured from tissue biopsies of each group; F1, Control without Cy; F2, Control; F3, 5 mg/kg/day TBBPA; F4, 1 mg/kg/day AMP. The experiment shown is representative of three studies in which similar differences were observed.

**Figure 4:**
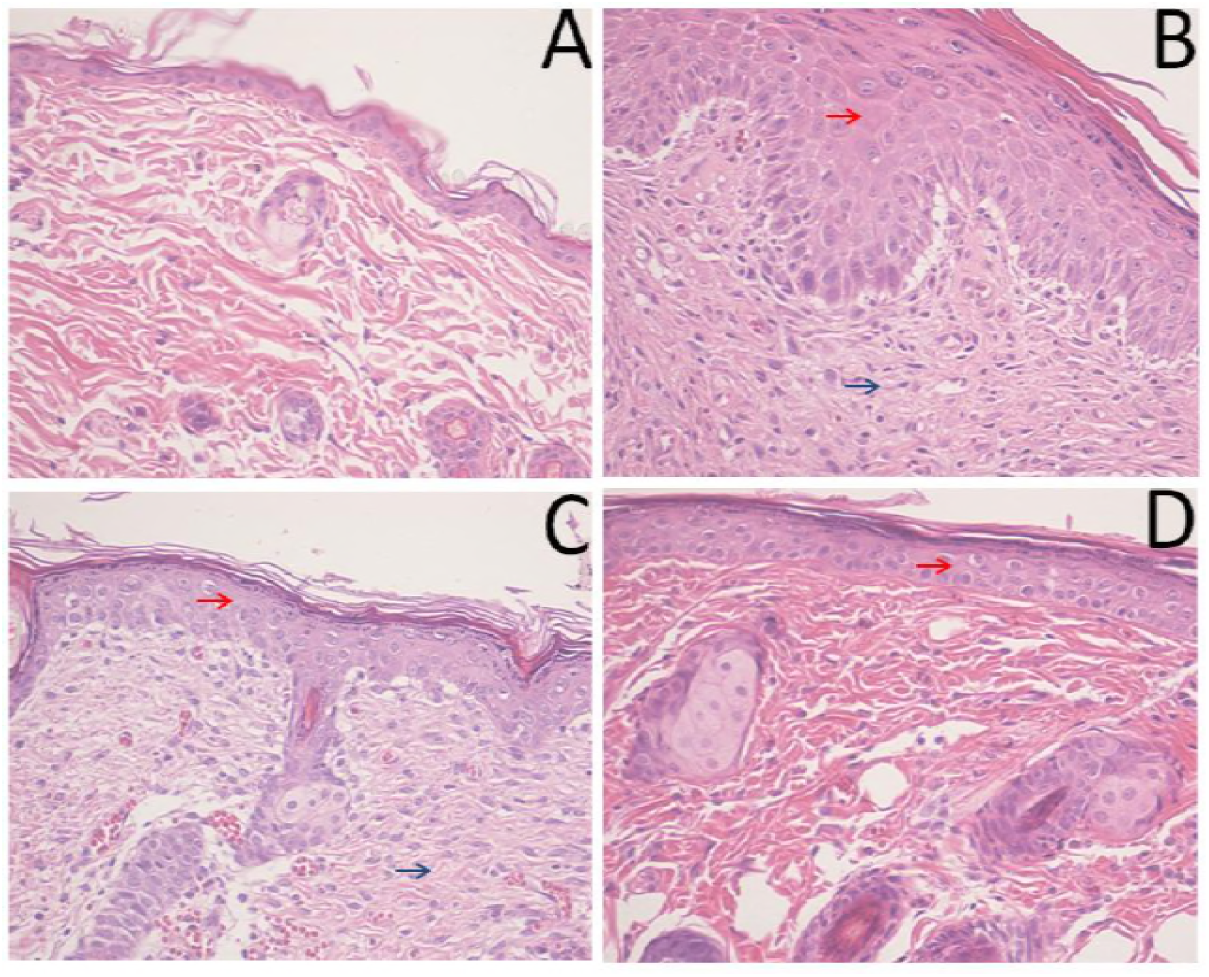
Histopathological examination of mouse wound surface, Hematoxylin-eosin staining, ×*200* magnification. A, Control without Cy; B, Control; C, 5 mg/kg/day TBBPA; D, 1 mg/kg/day AMP. Red arrows indicate thicker and tighter epidermis layer compared to the control group. Blue arrow indicate tighter dermis layer, inflammatory response and dissociative epidermis cells compared to control group (A).

The safety of TBBPA was first explored in mice. Upon 5 mg/kg TBBPA administration, no obvious adverse effects on body weight, feeding behavior, and physical activity of the mice were observed. This indicated that a low dose of TBBPA was not toxic and relatively safe. Acute dermal, oral and inhalation toxicity of TBBPA has been tested. The results showed that TBBPA exhibited low acute toxicity by multiple exposure routes (dermal LD_50_ > 10,000 mg/kg in rabbits, oral LD_50_ > 5,000 mg/kg in rats and inhalation LD_50_ > 10,000 mg/m^3^ in rats and mice). TBBPA does not cause skin, eye or respiratory irritation in humans or tested animals (12, 13). Chronic and sub-chronic toxicity studies of TBBPA show that it has a low hazard concern (13). Upon dietary exposure of TBBPA for 70 days in Wistar rats, no significant changes in reproduction were observed. TBBPA doses up to 3,000 mg/kg caused no histopathological changes in the organs assessed (14). In a two-generation toxicity study with doses of 0, 10, 100 or 1000 mg/kg/day TBBPA in Sprague Dawley rats, no obvious effects on reproduction, fertility, or developmental toxicity were observed. TBBPA shows low acute toxicity in mice, with an LD_50_ > 20 g/kg (15). With an NOAEL of 1,000 mg/kg, administration of 5 mg/kg TBBPA for 3 days should be relatively safe (13). No apparent physiological changes were observed in mice treated with 5 mg/kg TBBPA by gavage for 3 days (data not shown), adding evidence to the hypothesis that short-term exposure to a low dose of TBBPA should be safe.

This study evaluated the *in vivo* antimicrobial effects of TBBPA in a murine model of SA skin infection. We found that administration of TBBPA significantly reduced bacterial counts in the wound compared with vehicle, demonstrating the potential of this compound as a novel antimicrobial. The relative low MIC values of TBBPA suggest that this compound has potential as a candidate for the treatment of skin infections caused by SA *in vivo*. Ultimately, this compound may lead to specific therapeutic approaches to target SA infectious diseases. New compounds could be designed based on the structure of TBBPA, making modifications to improve the safety, bioavailability and treatment efficacy. Additional work is needed to further elucidate the specific antimicrobial mechanism of TBBPA and its application in skin infection therapy.

## Acknowledgements

This work was financially supported by grants from the Strategic Priority Research Program of the Chinese Academy of Sciences (XDB14030501) and the National Natural Science Foundation of China (21777061, 21477049, 21507155). We thank MD. Zhi Li from Wuhan General Hospital of Guangzhou Military for providing all the pathogenic strains and Dr. Song Hu from Jianghan University for his kind help in providing the MRSA strains. We thank Sarah Bubeck, Ph.D for editing the English text of a draft of this manuscript.

